## supplemental figures for "The subthalamic nucleus contributes causally to perceptual decision-making in monkeys"

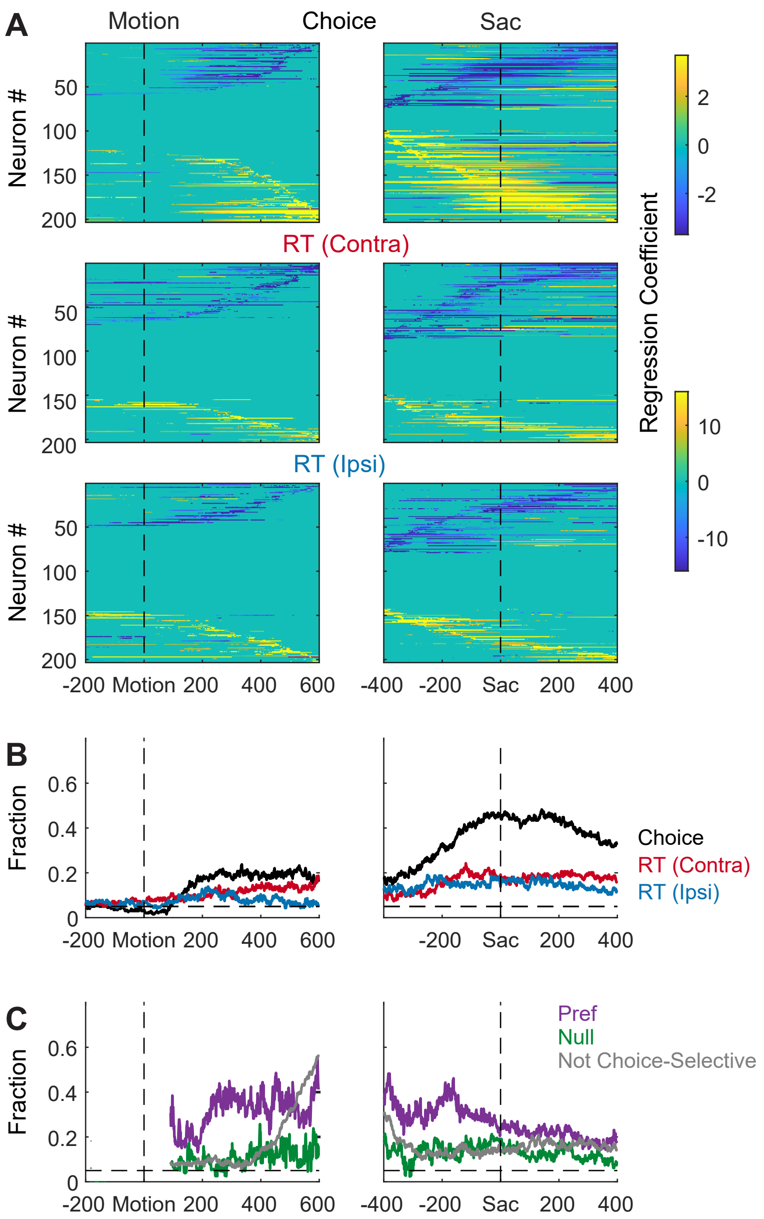


**Suppl. Figure 1. STN activity is modulated by choice and RT.** Same format as Figure 2, except using choice and RT as regressors.

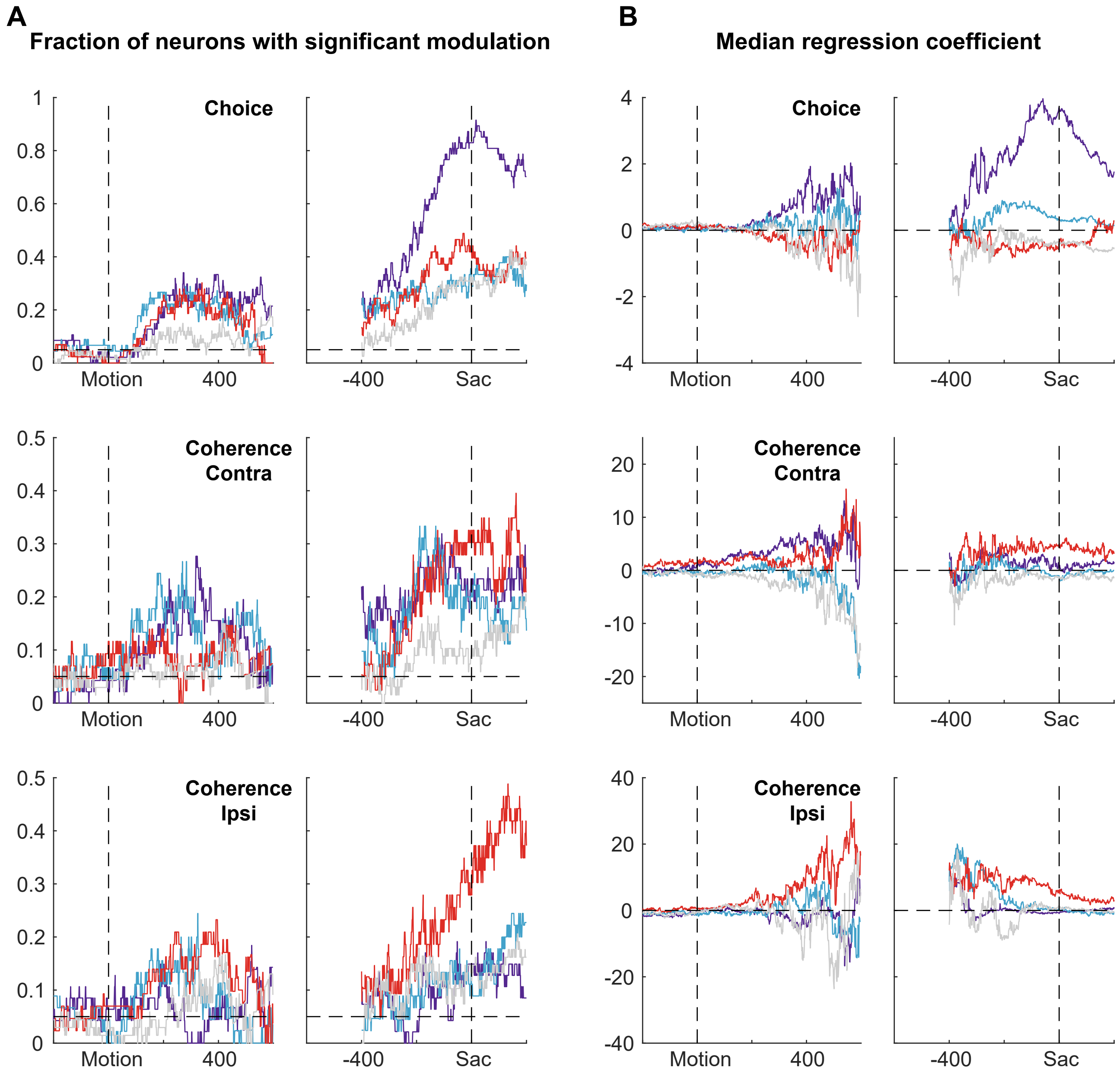


**Suppl. Figure 2. Summary of regression results, separated for different subpopulations.** A: fractions of neurons in each category that showed significant modulation (t-test, $p<0.05$) at each time window by choice (top), coherence for trials with contralateral choices (middle), and coherence for trials with ipsilateral choices (bottom). Dashed horizontal lines indicate the 5% chance level. Colors indicate categories as in Figure 3. B: median values of regression coefficients for choice and coherence as a function of time.

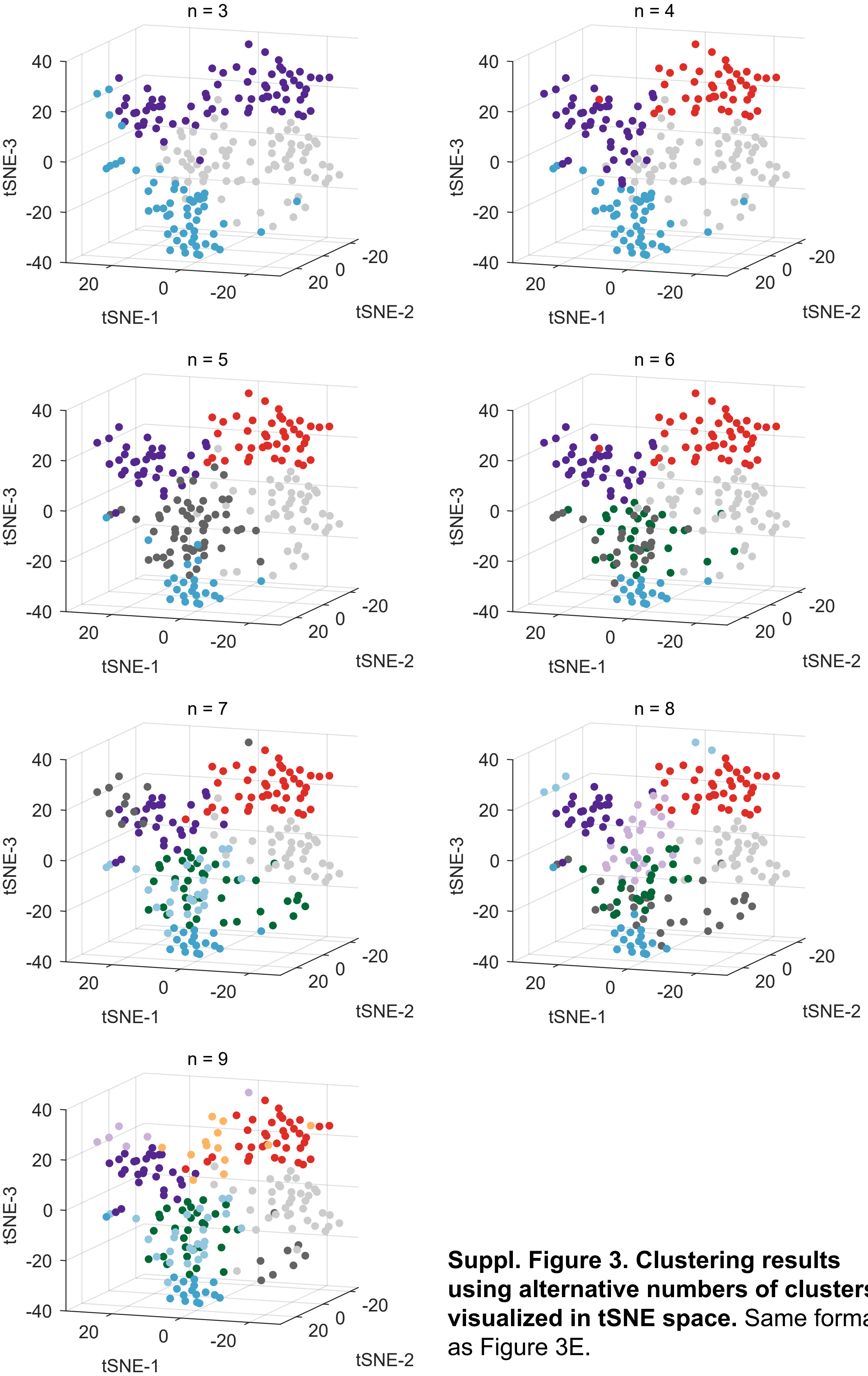


**Suppl. Figure 3. Clustering results using alternative numbers of clusters, visualized in tSNE space.** Same format as Figure 3E.

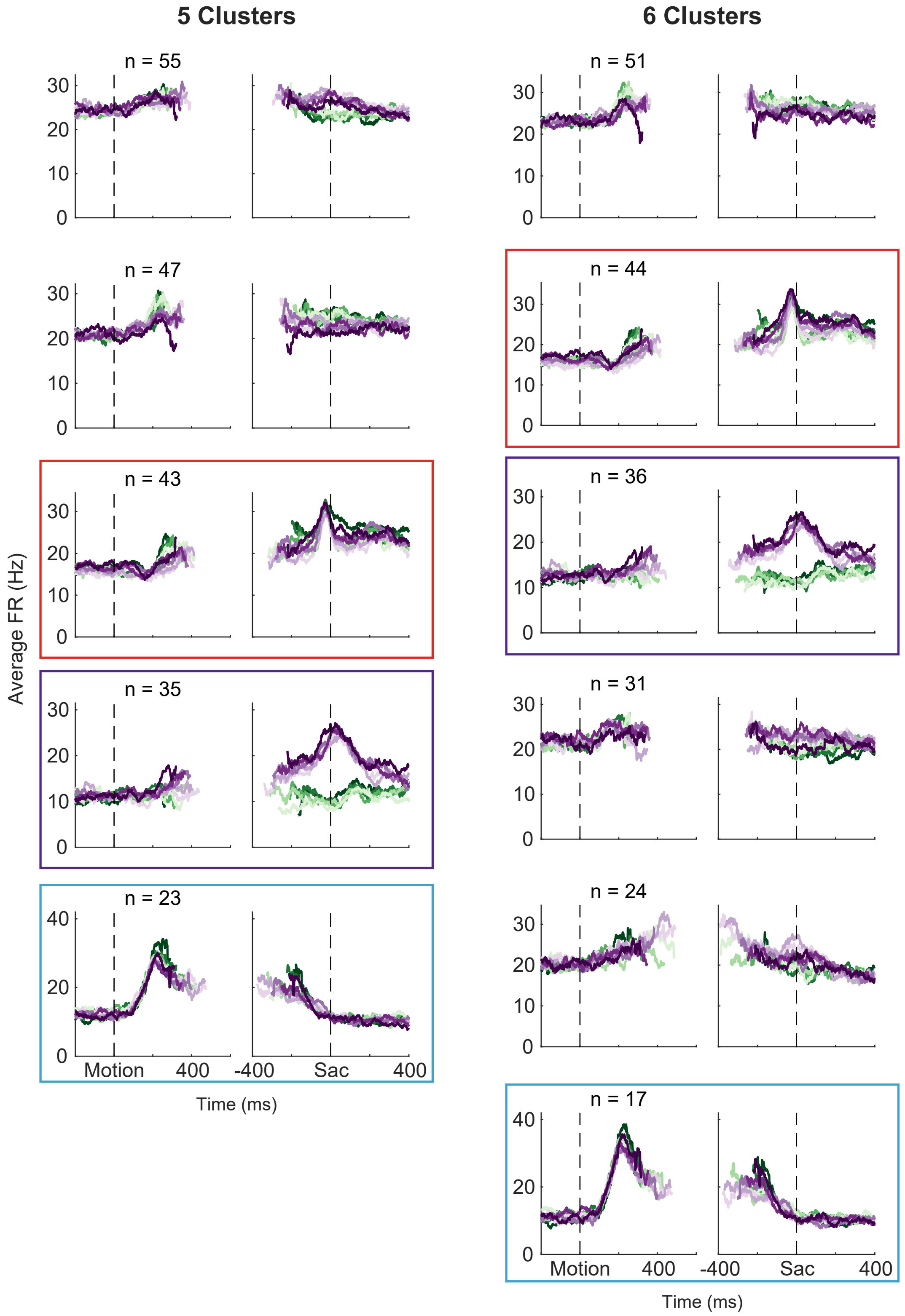


**Suppl. Figure 2. Clustering results using alternative numbers of clusters, visualized as average firing rates for each cluster.** Same format as Figure 3B and D. Colored boxes highlight the clusters corresponding to those identified in Figure 3.

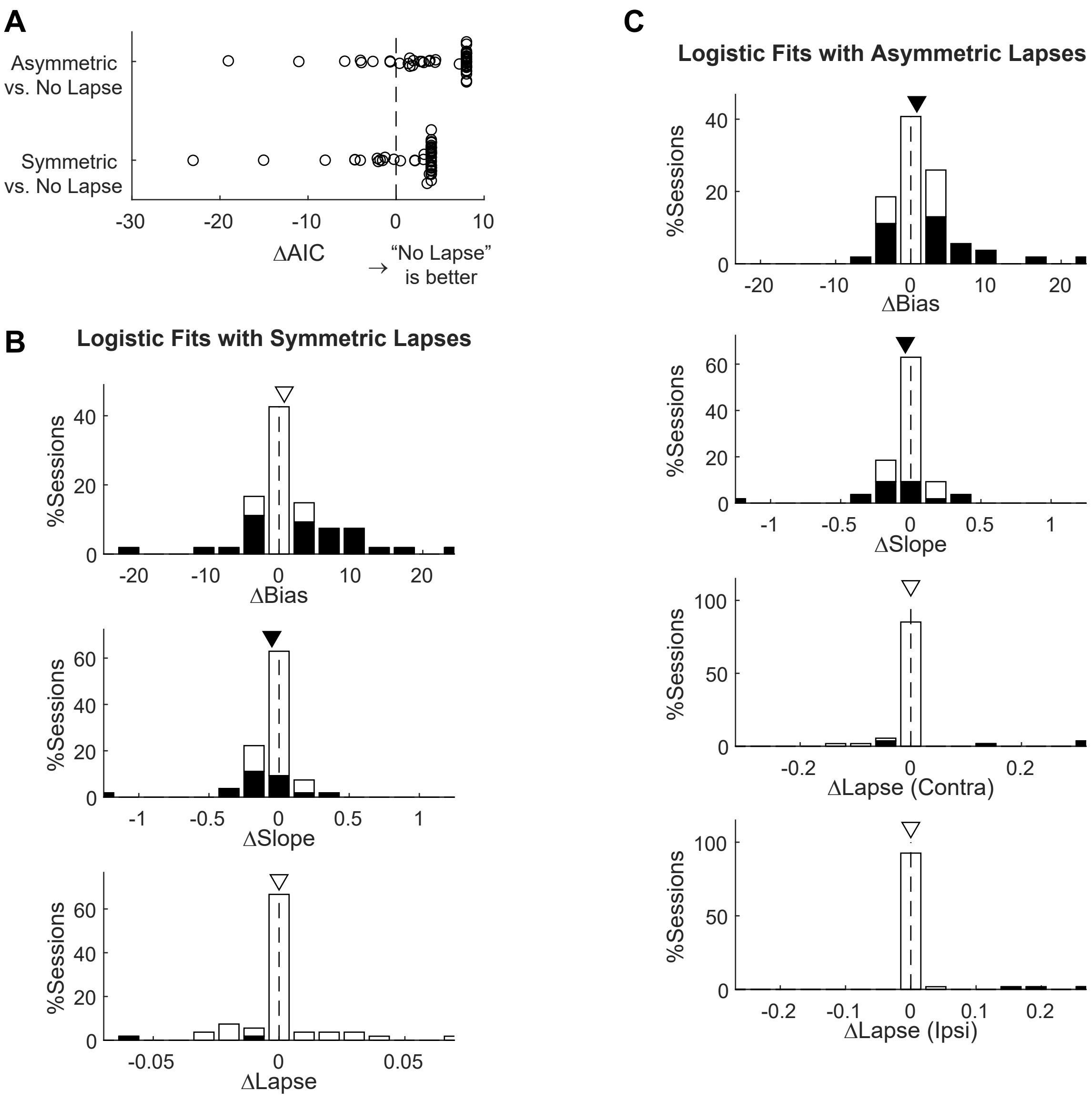


**Suppl Figure 5. Comparison of different logistic models**. A, The No Lapse model was associated with the lowest AIC for most sessions. The Symmetric Lapse model was associated with lower AICs for 12 sessions. The Asymmetric Lapse model was associated with lower AICs for 8 sessions. B, Histograms of microstimulation effects on bias, slope, and lapse terms in the Symmetric Lapse model. C, Histograms of microstimulation effects on bias, slope, and two lapse (for each choice) terms in the Asymmetric Lapse model. Same format as the histograms in Figure 5D.

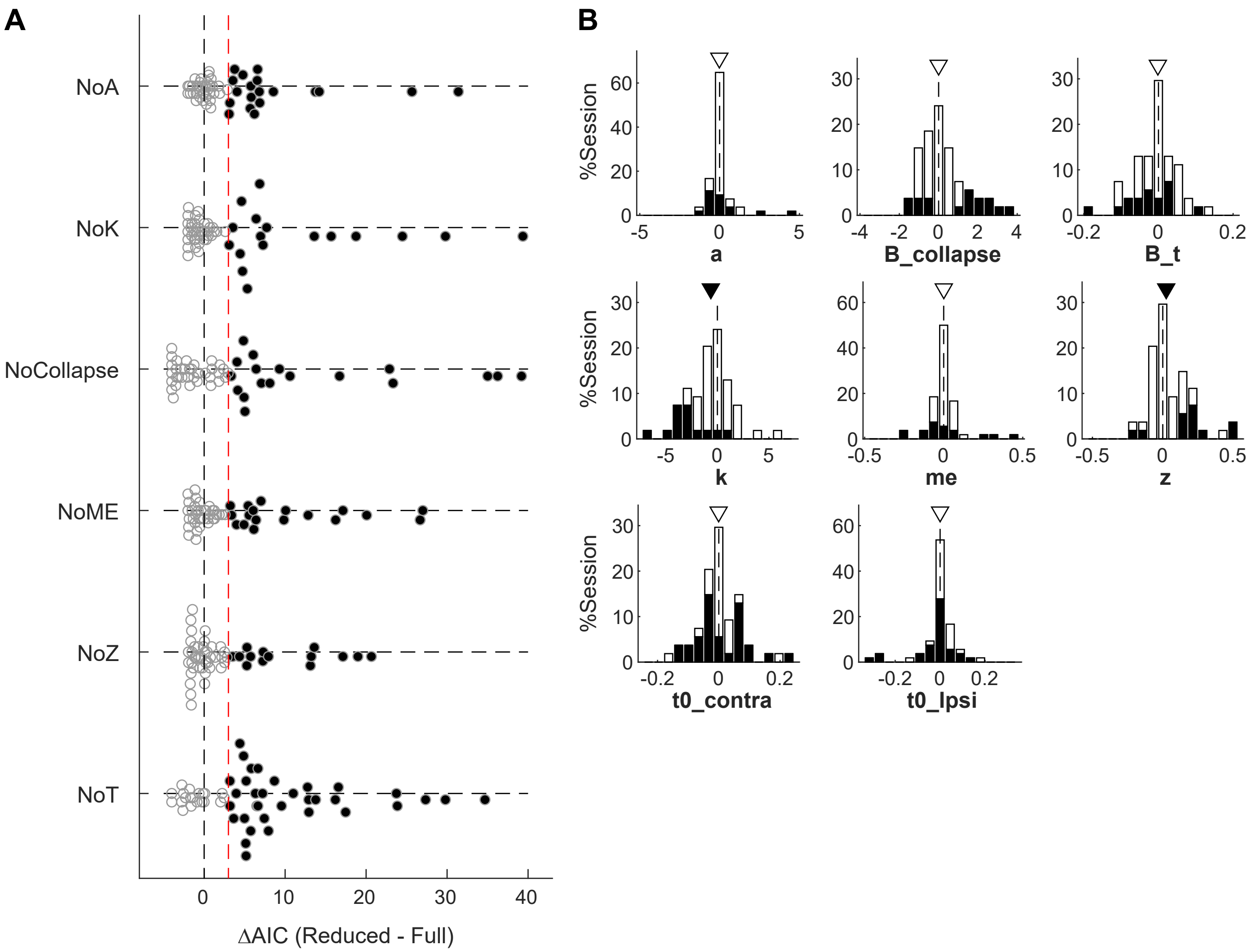


**Suppl Figure 6.** A, Differences in AIC between reduced and Full models. Filled circles indicate sessions for which $AIC_{Reduced} - AIC_{Full} > 3$ (red line). Note that for three sessions, the Full model outperformed the None model but not any of the reduced models. B, Histograms of difference in DDM parameters between trials with and without microstimulation. Filled bars represent sessions considered to show significant microstimulation effects on the given parameter, based on AIC comparisons. Triangles indicate median values. Filled triangles: Wilcoxon sign-rank test, $p < 0.05$.

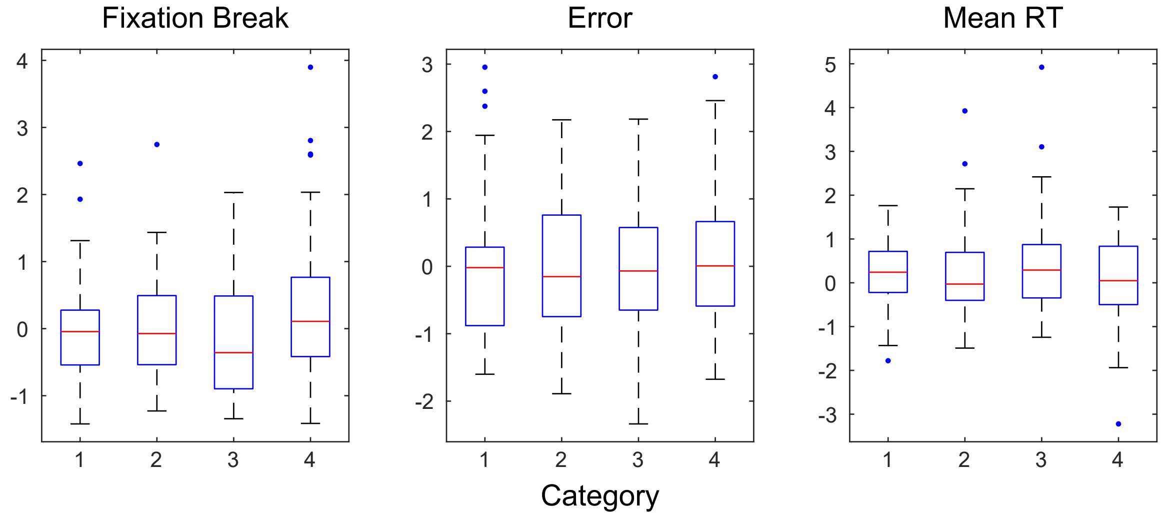


**Suppl. Figure 7. Indices of motivational state did not differ among sessions with different neuron subpopulations.** Panels show the summary of rate of fixation break (left), overall error rate (middle) and mean RT (right) for the four categories identified in Figure 3. All indices were z-scored across sessions for each monkey. Red lines indicate median values. The bottom and top edges of the box indicate the 25th and 75th percentiles, respectively. The whiskers extend to the most extreme data points not considered outliers, and the outliers are plotted individually as dots. ANOVA, $p = 0.06$, 0.91, and 0.29, respectively. No significant difference was observed for each monkey separately ($p > 0.08$ and 0.12 for all indices for monkeys C and F, respectively).
